# Smaller limbic structures are associated with greater immunosuppression in over 1000 HIV-infected adults across five continents: Findings from the ENIGMA-HIV Working Group

**DOI:** 10.1101/724583

**Authors:** Talia M. Nir, Jean-Paul Fouche, Jintanat Ananworanich, Beau M. Ances, Jasmina Boban, Bruce J. Brew, Linda Chang, Joga R. Chaganti, Christopher R.K. Ching, Lucette A. Cysique, Thomas Ernst, Joshua Faskowitz, Vikash Gupta, Jaroslaw Harezlak, Jodi M. Heaps-Woodruff, Charles H. Hinkin, Jacqueline Hoare, John A. Joska, Kalpana J. Kallianpur, Taylor Kuhn, Hei Y. Lam, Meng Law, Christine Lebrun-Frenay, Andrew J. Levine, Lydiane Mondot, Beau K. Nakamoto, Bradford A. Navia, Xavier Pennec, Eric C. Porges, Cecilia M. Shikuma, April D. Thames, Victor Valcour, Matteo Vassallo, Adam J. Woods, Paul M. Thompson, Ronald A. Cohen, Robert Paul, Dan J. Stein, Neda Jahanshad, for the ENIGMA-HIV Working Group

**Author notes:** Corresponding Author: Neda Jahanshad, PhD, Assistant Professor of Neurology and Biomedical Engineering, Imaging Genetics Center, Mark and Mary Stevens Neuroimaging and Informatics Institute, Keck School of Medicine of USC, University of Southern California, 4676 Admiralty Way, Marina del Rey, CA 90292, USA;,; Phone: 323-44-BRAIN. These authors jointly supervised this work (Ronald A. Cohen, Robert Paul, Dan J. Stein, Neda Jahanshad).

## Abstract

**Background:** Human immunodeficiency virus type-1 (HIV) infection can be controlled with combination antiretroviral therapy (cART), but neurocognitive impairment remains common even in chronic and treated HIV-infected (HIV+) cohorts. Identifying the neuroanatomical pathways associated with infection has the potential to delineate novel neuropathological processes underlying persisting deficits, yet individual neuroimaging studies have yielded inconsistent findings. The ENIGMA-HIV Working Group was established to harmonize data from diverse studies to identify the common effects of HIV-infection on brain structure.

**Methods:** Data were pooled from 12 independent neuroHIV studies from Africa, Asia, Australia, Europe, and North America. Volume estimates for eight subcortical brain regions were extracted from T1-weighted MRI from 1,044 HIV+ adults (aged 22-81 years; 72.4% on cART; 70.3% male; 41.6% with detectable viral load (dVL)), to identify associations with plasma markers reflecting current immunosuppression (CD4+ T-cell count) or dVL. Follow-up analyses stratified data by cART status and sex. Bonferroni correction was used to determine statistical significance.

**Findings:** Lower current CD4+ count was associated with smaller hippocampal (*β* = 20.3 mm^3^ per 100 cells/mm^3^; *p* = 0.0001) and thalamic volumes (*β* = 29.3; *p* = 0.003); in the subset of participants not on cART, it was associated with smaller putamen volumes (*β* = 65.1; *p* = 0.0009). On average, a dVL was associated with smaller hippocampal (Cohen’s *d* = 0.24; *p* = 0.0003) and amygdala volumes (*d* = 0.18; *p* = 0.0058).

**Interpretation:** In HIV+ individuals across five continents, smaller limbic volumes were consistently associated with current plasma markers. As we assessed cohorts with different inclusion/exclusion criteria and demographic distributions, these deficits may represent a generalizable brain-signature of HIV infection in the cART era. Our findings support the importance of achieving viral suppression and immune restoration for maintaining brain health.

**Funding:** This work was supported, in part, by NIH grant U54 EB020403.

**Research in Context:** *Evidence before this study:* HIV type-1 infection can be managed with antiretroviral therapy, however neurocognitive impairment persists even in treated HIV+ individuals. Given the challenges associated with standardized cognitive testing, there is a need to identify quantitative markers of central nervous system impairment. A number of neuroimaging studies have reported brain abnormalities in HIV-infected patients; however, prior studies investigating associations between CD4+ T-cell count or HIV viral load and subcortical brain volume report variable effect sizes and regional distributions of effects, limiting the generalizability of the conclusions drawn to date. We have conducted a literature search for reports in English language journals published until June 2019, using the following search terms: HIV AND subcortical AND neuroimaging AND brain AND viral load AND RNA AND CD4. After removing studies that were not applicable, there were 30 studies investigating CD4+ T-cell count and viral load associations with subcortical brain structure.

*Added value of the study:* The aim of the current study was to investigate structural brain associations with two biomarkers universally used to monitor immune function and treatment response, namely plasma RNA viral load and CD4+ T-cell counts. Prior analyses have been performed in smaller, heterogeneous cohorts, but by combining data across cohorts, we can identify consistent associations between brain volume and indicators of HIV infection across cohorts. The ENIGMA-HIV Working Group was established to identify common neurobiological signatures of the HIV-infected brain by harmonizing data analysis from HIV neuroimaging studies worldwide. The value of this dataset is that it is well-powered and representative of many HIV+ people living in the cART era.

*Implications of all the available evidence:* Our results provide robust evidence that despite demographic and clinical heterogeneity among HIV-infected individuals, brain abnormalities are consistently linked to HIV viral load and immunosuppression. This supports the importance of achieving viral suppression and immune system restoration in maintaining brain health in people living with HIV. The vulnerability of limbic regions, found in this study, extends beyond the classically implicated regions of the basal ganglia; this suggests that these regions remain an important target of cART era HIV research, especially given their heightened vulnerability to age-associated atrophy and neurodegeneration.

## Introduction

In the era of globally accessible combination antiretroviral therapy (cART), morbidity and mortality rates have dramatically decreased for individuals who have contracted the human immunodeficiency virus type-1 (HIV). HIV-infected (HIV+) individuals on cART are now expected to reach near-normal lifespans. However, HIV-related comorbidities, including symptoms of brain dysfunction, remain common even in treated HIV+ individuals.^1,2^

The prevalence of neurocognitive impairment (NCI) in HIV+ populations varies across studies: some reports suggest that cognitive difficulties are present in nearly half of HIV+ individuals, but other studies report less than 20% are affected.^3^ This variability may reflect heterogeneous viral-host dynamics and cART treatment status. Other factors include inconsistencies in educational attainment or quality, and neuropsychological testing methods, including the use of appropriate normative data and how neurological and psychiatric confounds were handled. In studies using standard neuropsychological assessments and appropriate normative data, there is robust evidence that even mild NCI can interfere with the capacity to perform instrumental activities of daily living (IADLs).^2^

Plasma viral load (VL) and CD4+ T-cell count are two biomarkers universally used to monitor immune function and treatment response. These are also the most consistently available clinical markers in human studies of HIV, but the degree to which they capture CNS impairment is not fully understood. Low nadir CD4+ has been identified as a predictor of NCI in the cART era,^4^ suggesting that severe immunosuppression may lead to persistent and potentially irreversible brain injury. However, nadir CD4+ is frequently self-reported, so it may be unreliable or unknown. The duration of immunosuppression before recovery and the highest VL may also not be well documented. Immune restoration and viral suppression are tracked through routine clinical assessments, and may be achieved through modifiable factors including timely HIV testing, as well as treatment initiation and compliance. The neurological implications of maintaining or achieving healthy targets, are therefore important to establish.

Structural neuroimaging provides a promising array of quantitative biomarkers for assessing CNS function and decline. Neuroimaging biomarkers have provided important non-invasive means to understand many psychiatric and neurodegenerative diseases, and to monitor efficacy of clinical interventions.^5^ Neuroimaging studies of HIV+ individuals tend to show abnormal volumes of subcortical structures in infected individuals,^6^ and associations between such structural changes and NCI;^7^ however, inconsistencies in the effect sizes and regional distribution of brain abnormalities associated with CD4+ count or VL in HIV+ individuals have limited the generalizability of the conclusions drawn to date. These inconsistent findings are likely driven in part by differences in demographic characteristics such as age and sex, socioeconomic status, lifestyle factors, history of trauma, substance use, and clinical heterogeneity of study participants, including viral suppression, cART timing/adherence, and other comorbidities. For instance, a study of untreated, chronically infected individuals reported a positive association between subcortical volumes and current CD4+ T-cell counts,^8^ while another study evaluating a diverse sample reported negative associations.^9^ Yet effects are not necessarily population specific, as studies focusing on individuals on long-term treatment^10^ and those who are cART naive^11^ have both reported no associations with CD4+ count. Different findings across studies may also reflect methodological heterogeneity, including differences in 1) statistical power with variable sample sizes, in addition to unequal representation of important HIV disease factors and NCI prevalence; 2) MRI scanners and image acquisition protocols; and 3) image processing or statistical analysis techniques. By reducing methodological variability in image processing, boosting sample size, and assessing a diverse set of HIV+ cohorts, a generalizable pattern of HIV-related brain effects may be identified.

The HIV Working Group was established within the Enhancing Neuro Imaging Genetics through Meta Analysis (ENIGMA) consortium to pool data from neuroimaging studies using harmonized data analysis pipelines. The working group is a growing international collaboration open to all researchers investigating the neurological consequences of HIV infection. This current study included active involvement from investigators from 12 neuroHIV studies from six different countries: the United States, France, Serbia, Australia, Thailand, and South Africa (**Table 1**). We aimed to investigate structural brain volume associations with the most commonly collected clinical assessments of HIV burden; we surveyed CD4+ T-cell counts and the detectability of viral RNA, in plasma, and determined their relationships with MRI-derived subcortical brain volumes in HIV+ individuals. The strength of this large dataset is that it is representative of many HIV+ individuals currently living in the cART era (including those who are not treated, with and without viral suppression). Subcortical regions remain a target of HIV infection, therefore, these nuclei represent an ideal benchmark against which the degree of immune recovery, viral suppression, and cART use may be assessed.

**Table 1.** Demographic and clinical information by study and scanning site.

| Site name | Total N | Male % (N) | Age [yrs] Mean (SD), Range | on cART % (N) | Detectable Viral Load % (N) | CD4+ Count cells/mm<sup>3</sup> Mean (SD) | CD4+ Count < 200 cells/mm<sup>3</sup> % (N) | Viral Load > 400 copies/mL % (N) |
| --- | --- | --- | --- | --- | --- | --- | --- | --- |
| **HIVNC Consortium (7 Sites), United States** | 229 | 85.2 (195) | 48.6 (8.3) 24-71 | 100 (229) | 27.4 (62) N=226 | 374.0 (229.6) | 22.3 (51) | 17.7 (40) N=226 |
| Site 1:<br>University of California, San Diego | 25 | 96.0 (24) | 47.3 (4.6) 37-56 | 100 (25) | 59.1 (13) N=22 | 308.8 (306.7) | 48.0 (12) | 31.8 (7) N=22 |
| Site 2:<br>Harbor UCLA Medical Center | 54 | 79.6 (43) | 46.6 (8.9) 24-70 | 100 (54) | 14.8 (8) | 350.6 (174.4) | 27.8 (15) | 9.3 (5) |
| Site 3:<br>Stanford University | 10 | 90.0 (9) | 46.4 (10.4) 31-62 | 100 (10) | 0 (0) | 313.5 (157.5) | 20.0 (2) | 0 (0) |
| Site 4:<br>Colorado | 37 | 97.3 (36) | 49.5 (7.8) 31-66 | 100 (37) | 37.8 (14) | 439.4 (241.8) | 10.8 (4) | 16.2 (6) |
| Site 5:<br>Pittsburgh | 19 | 94.7 (18) | 49.9 (9.7) 34-71 | 100 (19) | 21.1 (4) | 438.9 (298.2) | 15.8 (3) | 15.8 (3) |
| Site 6:<br>Rochester University | 42 | 64.3 (27) | 48.6 (7.8) 26-62 | 100 (42) | 26.2 (11) | 379.3 (206.0) | 14.3 (6) | 16.7 (7) |
| Site 7:<br>University of California, Los Angeles | 42 | 90.5 (38) | 50.7 (8.3) 28-69 | 100 (42) | 28.6 (12) | 364.9 (224.3) | 21.4 (9) | 28.6 (12) |
| University of Hawaii, United States | 53 | 84.9 (45) | 50.9 (8.0) 40-71 | 100 (53) | 11.3 (6) | 491.1 (208.5) | 7.5 (4) | 3.8 (2) |
| University of California, San Francisco, United States | 50 | 98.0 (49) | 63.6 (2.5) 60-69 | 98 (49) | 28.6 (14) N=49 | 529.3 (218.5) | 0 (0) | 8.2 (4) N=49 |
| Brown University, United States | 79 | 60.8 (48) | 45.2 (9.5) 23-65 | 83.5 (66) | 27.8 (22) | 476.2 (228.8) | 8.9 (7) | 26.6 (21) |
| University of California, Los Angeles, United States (Hinkin) | 12 | 100 (12) | 46.2 (8.5) 26-57 | 75 (9) | 50.0 (6) | 604.1 (289.1) | 8.3 (1) | 33.3 (4) |
| University of California, Los Angeles, United States (Thames) | 54 | 90.7 (49) | 50.5 (12.9) 24-76 | 100 (54) | 42.6 (23) | 613.7 (284.5) | 3.7 (2) | 18.5 (10) |
| University of New South Wales, Australia (Brew) | 41 | 97.6 (40) | 52.8 (8.1) 39-75 | 100 (41) | 29.3 (12) | 596.8 (271.5) | 0 (0) | 19.5 (8) |
| University of New South Wales, Australia (Cysique) | 68 | 100 (68) | 55.3 (6.7) 44-69 | 100 (68) | 0.01 (1) | 549.46 (273.63) | 7.4 (5) | 0.01 (1) |
| SEARCH 011 Consortium, Thailand | 61 | 42.6 (26) | 34.2 (7.0) 22-56 | 0 (0) | 100 (61) | 236.0 (139.0) | 41.0 (25) | 100 (61) |
| University of Cape Town, South Africa | 181 | 13.8 (25) | 32.4 (5.0) 22-46 | 0 (0) | 97.4 (148) N=152 | 225.5 (149.2) | 52.5 (95) | 87.8 (129) N=147 |
| Nice University, France | 155 | 78.7 (122) | 45.4 (10.0) 22-81 | 75.9 (126) | 36.1 (56) | 580.9 (277.6) | 7.1 (11) | 25.2 (39) |
| University of Novi Sad, Serbia | 61 | 90.2 (55) | 44.3 (10.9) 25-66 | 100 (61) | 16.4 (10) | 616.3 (346.2) | 1.5 (1) | 16.4 (10) |

## Methods

### Participants and Clinical Assessments

T1-weighted magnetic resonance imaging (MRI) brain scans and clinical data from 1,044 HIV+ adult participants (aged 22-81 years; 70.3% male) were collected at 18 sites from 12 independent studies. The participating studies include the multisite HIVNC Consortium (N=229 scanned across 7 sites), Brown University (N=79), University of Hawaii (N=53), University of California San Francisco (N=50) and two groups from the University of California Los Angeles (Hinkin N=12; Thames N=54), all in the United States; Nice University Hospital in France (N=155); the University of Cape Town in South Africa (N=181); two groups from the University of South Wales in Australia (Brew N=41; Cysique N=68); a group from Serbia (N=61), and the SEARCH-011 study from Thailand (N=61). Inclusion and exclusion criteria for each study are summarized in **Supplementary Table 1.** Clinical assessments at the time of scan included current CD4+ T-cell counts (cells/mm^3^) and HIV plasma RNA VL (copies/mL).

Participant demographic and clinical characteristics for each of the 12 studies are reported in **Table 1**. Two transgender participants were excluded. Each study obtained approval from their local ethics committee or institutional review board; participants signed an informed consent form at each participating site.

### Image acquisition, processing, and quality assurance

T1-weighted brain MRI scans were acquired at each site, on either a 3 or 1.5 tesla scanner platform. Acquisition protocols are detailed in **Supplementary Table 2**. Intracranial volume estimation and regional segmentations were conducted using FreeSurfer version 5.3. Volumes were extracted from eight regions of interest (ROIs): thalamus, caudate, putamen, pallidum, hippocampus, amygdala, nucleus accumbens, and lateral ventricles. Volume extraction and quality control were completed using publically available protocols (http://enigma.usc.edu/protocols/). The average of the left and right volumes for each ROI was evaluated. Histograms were also created from each site’s data to investigate normality of the data distribution for each ROI, and statistical outliers were identified. If the mean of an individual’s subcortical volume was more than 3 standard deviations from the mean for the site, it was flagged for a more extensive quality control and possible removal from the analysis. The 1,044 scans included in the overall study represent only the scans for which all segmentations were of sufficient quality. Quality assurance also included a feasibility study to ensure the pooled sample was able to capture known negative associations between brain volume and age (**Supplementary Methods and Results**).

### Statistical analysis

#### Associations between brain volumes and HIV plasma markers

Random effects multiple linear regressions were performed to evaluate associations between each of the eight regional brain volumes and 1) current CD4+ T-cell count (cells/mm^3^) or 2) a binary variable indicating a detectable (1) or undetectable (0) VL. For each participant, VL was determined to be detectable or undetectable according to the detection threshold at the respective collection site, which varied (range: 2-400 copies/mL, please see **Supplementary Table 3**). To account for differences in MRI scanner and acquisition, MRI collection site was used as the random-effects grouping variable; fixed-effects covariates included age, sex, the interaction between age and sex, and estimated total intracranial volume (eICV) to adjust for variability in head size. See **Supplemental Methods and Results** for details. Effect sizes, after accounting for all covariates, were estimated as Cohen’s *d*-values for dichotomous factors and *r*-values (partial correlation coefficients) for continuous variables. Bonferroni correction was used to control the family-wise error rate at 5%; *p*-value was significant if ≤ 0.0063 = 0.05/8 (brain regions).

#### Stratification by cART status and by sex

*Post-hoc* analyses tested for associations between brain volumes and HIV plasma markers when the data were stratified by 1) cART status at the time of scan (cART+ or cART-), and 2) sex. The same analytic framework was used, removing sex and sex-by-age interactions from statistical models when stratifying by sex. Group differences in demographic and clinical factors were evaluated with chi-square tests for dichotomous factors, and two-tailed *t*-tests for continuous measures.

#### Validation analyses comparing harmonized thresholds, and meta-analyses

Three sets of validation analyses were performed: 1) dichotomizing CD4+ count based on the AIDS-defining threshold of 200 cells/mm^3^; 2) defining a common detectable VL (dVL) threshold across sites (400 copies/mL, the highest site-specific assay detection threshold used across all collection sites); and 3) comparing primary pooled statistical analyses to the meta-analytic framework used in other ENIGMA studies,^12^ where associations were conducted in each site separately, then aggregated using an inverse-variance weighted meta-analysis (see **Supplementary Methods and Results** for details).

## Results

Demographic and clinical characteristics of the entire ENIGMA-HIV cohort are summarized in **Table 2**. Compared to cART+ participants (N=756), cART-participants (N=288) were younger, had lower CD4+ counts, and a greater percentage had dVL. Similarly, compared to males (N=734), females (N=310) were younger, had lower CD4+ counts, and a greater percentage had dVL. Proportionally fewer females were on cART at the time of scan. Across participants, older age was associated with smaller volumes of all subcortical structures and larger ventricular volumes, as expected; results from the feasibility analysis of age are presented in **Supplementary Tables 4 and 5** and **Supplementary Figures 1 and 2**.

**Table 2.** Summary of demographic, clinical, and neuroanatomical information aggregated across all 12 participating studies of HIV+ adults, and stratified by cART status and by sex. Subcortical volume comparisons in this table are absolute and were not corrected for eICV, as was done in the formal analyses.

|  | Total | cART+ | cART- | cART+ vs<br>cART-<br>*P-Value | Male | Female | Male vs<br>Female<br>*P-Value |
| --- | --- | --- | --- | --- | --- | --- | --- |
| **Total N** | 1,044 | 756 (72.41%) | 288 (27.59%) |  | 734 (70.3%) | 310 (29.7%) |  |
| **Mean Age**<br>**yrs (SD), Range** | 45.46 (11.60),<br>22-81 | 49.71 (9.91)<br>23-81 | 34.29 (7.62)<br>22-66 | **4.93x10<sup>-108</sup>** | 48.44 (10.80),<br>22-81 | 38.40 (10.30),<br>22-70 | **1.16x10<sup>-39</sup>** |
| **% on**<br>**cART (N)** | 72.41% (756) | 100% (756) | -- | -- | 87.87% (645) | 35.81% (111) | **2.71x10<sup>-66</sup>** |
| **% with Detectable**<br>**Viral Load (N)** | 41.64% (421)<br>N=1011 | 22.21% (167)<br>N=752 | 98.07% (254)<br>N=259 | **3.20x10<sup>-101</sup>** | 31.68% (230)<br>N=726 | 67.02% (191)<br>N=285 | **2.28x10<sup>-25</sup>** |
| **% with Viral Load**<br>**> 400 copies/mL (N)** | 32.70% (329)<br>N=1006 | 12.63% (95)<br>N=752 | 92.13% (234)<br>N=254 | **1.40x10<sup>-120</sup>** | 21.55% (156)<br>N=724 | 61.35% (173)<br>N=282 | **1.25x10<sup>-33</sup>** |
| **Mean CD4+ Count**<br>**cells/mm<sup>3</sup> (SD)** | 441.40 (275.82) | 501.30 (276.05) | 284.26 (204.30) | **1.01x10<sup>-38</sup>** | 483.77 (278.40) | 341.07 (242.00) | **5.19x10<sup>-16</sup>** |
| **% with CD4+ Count**<br>**< 200 cells/mm<sup>3</sup> (N)** | 19.35% (202) | 10.71% (81) | 42.01% (121) | **2.57x10<sup>-30</sup>** | 13.76% (101) | 32.58% (101) | **2.01x10<sup>-12</sup>** |
| **Mean Thalamus**<br>**Volume mm<sup>3</sup> (SD)** | 7043.5 (979.5) | 7189.2 (978.6) | 6661.1 (874.4) | **2.73x10<sup>-16</sup>** | 7277.7 (963.4) | 6489.0 (773.2) | **2.44x10<sup>-39</sup>** |
| **Mean Caudate**<br>**Volume mm<sup>3</sup> (SD)** | 3587.3 (498.0) | 3616.1 (502.8) | 3511.8 (477.8) | **0.0020** | 3652.4 (500.1) | 3433.2 (458.3) | **1.57x10<sup>-11</sup>** |
| **Mean Putamen**<br>**Volume mm<sup>3</sup> (SD)** | 5206.6 (740.0) | 5183.3 (779.3) | 5267.8 (622.2) | **0.069** | 5270.0 (777.5) | 5056.51 (617.99) | **2.98x10<sup>-6</sup>** |
| **Mean Pallidum**<br>**Volume mm<sup>3</sup> (SD)** | 1517.5 (251.7) | 1478.6 (249.4) | 1619.6 (228.4) | **4.14x10<sup>-17</sup>** | 1517.9 (251.8) | 1516.6 (251.9) | 0.94 |
| **Mean Hippocampus**<br>**Volume mm<sup>3</sup> (SD)** | 4002.1 (544.9) | 4109.7 (498.0) | 3719.6 (562.2) | **1.11x10<sup>-22</sup>** | 4152.0 (507.5) | 3647.1 (459.8) | **2.97x10<sup>-47</sup>** |
| **Mean Amygdala**<br>**Volume mm<sup>3</sup> (SD)** | 152.8 (299.8) | 1620.1 (299.7) | 1376.2 (217.3) | **6.12x10<sup>-42</sup>** | 1640.9 (287.3) | 1344.2 (213.5) | **4.37x10<sup>-63</sup>** |
| **Mean Accumbens**<br>**Volume mm<sup>3</sup> (SD)** | 541.0 (128.2) | 527.4 (135.5) | 576.8 (98.3) | **1.66x10<sup>-10</sup>** | 539.8 (135.8) | 543.9 (108.1) | 0.60 |
| **Mean Lateral**<br>**Ventricles**<br>**Volume mm<sup>3</sup> (SD)** | 19844.6 (14156.4) | 21935.1 (14969.2) | 14357.0 (9853.4) | **2.07x10<sup>-20</sup>** | 22204.9 (15009.2) | 14256.2 (9873.2) | **1.12x10<sup>-22</sup>** |

\*Differences in dichotomous factors were assessed using chi-squared tests; for continuous measures, differences were assessed using two-tailed *t*-tests.

### Associations between brain volumes and HIV plasma measures

Across all participants (N=1,044), lower CD4+ counts were associated with smaller hippocampal (*r* = 0.12; *p* = 0.0001; **Table 3**) and smaller thalamic volumes (*r* = 0.09; *p* = 0.003). dVL (N=1,011) was associated with smaller hippocampal (Cohen’s *d* = -0.24; *p* = 0.0003; **Table 4**) and amygdala volumes (*d* = -0.18; *p* = 0.0058). Associations between CD4+ counts or dVL and brain volumes remained consistent after additionally adjusting for cART status in the statistical models. To ensure both CD4+ count and dVL were independently associated with hippocampal volume, we included both predictors in the same model and found both remained significant (dVL: *d* = -0.20, *p* = 0.0023; CD4+: *r* = 0.11; *p* = 0.0008).

**Table 3.**
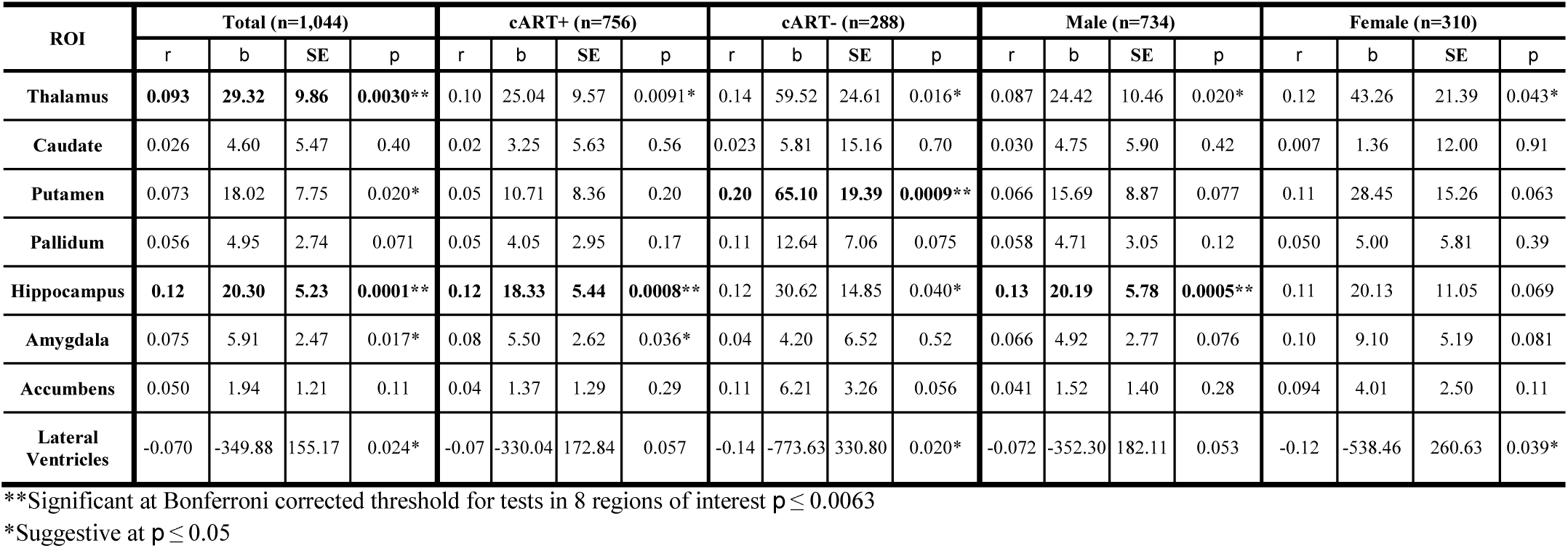
*R*-values (partial correlation coefficients), *b*-values (unstandardized regression slopes reflecting change in volume (mm^3^) for every 100 cells/mm^3^ change in CD4+ count), standard errors (SE), and *p*-values from associations between regional brain volumes and CD4+ count at the time of scan across all 1,044 HIV+ participants, and separately in the subset of 756 cART+ participants, 288 cART-participants, 734 males, and 310 females.

### Stratification by cART status and by sex

**Table 3** (CD4+) and **Table 4** (dVL) detail regional findings within stratified groups. In the subset of cART+ participants, smaller hippocampal volumes were associated with lower CD4+ count and dVL (CD4+: *r* = 0.12; *p* = 0.0008; dVL: *d* = -0.31, *p* = 0.0005). cART+ participants also showed amygdala volume associations with dVL (*d* = -0.30; *p* = 0.0005). In cART-participants, smaller putamen volumes were associated with lower CD4+ counts (*r* = 0.20; *p* = 0.0009); as only 5 cART-participants had an undetectable VL, dVL was not assessed. Associations in males mirrored those found in the cART+ subgroup: hippocampal volumes were associated with both CD4+ and dVL (CD4+: *r* = 0.13; *p* = 0.0005; dVL: *d* = -0.29; *p* = 0.0004), and amygdala volumes were associated with dVL (*d* = -0.25; *p* = 0.0024). We found no statistically significant associations between plasma measures and any brain volume in females; however we found a similar pattern in the ranking of regional effect sizes between males and females (CD4+: Pearson’s *r* = 0.91; *p* = 0.002; dVL: *r* = 0.79; *p* = 0.02; **Supplementary Figure 3**).

**Table 4.** Cohen’s *d* effect sizes, standard errors (SE), and *p*-values for associations between regional brain volumes and detectable VL at the time of scan across all 1,011 HIV+ participants with VL information, and separately in the subset of 752 cART+ participants, 726 males, and 285 females. We did not assess VL in those off treatment due to the limited number of individuals in this subgroup with undetectable VL (n=5).

| ROI | Total (n=1011) |  |  | cART+ (n=752) |  |  | cART- (n=259) |  |  | Male (n=726) |  |  | Female (n=285) |  |  |
| --- | --- | --- | --- | --- | --- | --- | --- | --- | --- | --- | --- | --- | --- | --- | --- |
|  | Cohen's *d* | SE | *p* | Cohen's *d* | SE | *p* | Cohen's *d* | SE | *p* | Cohen's *d* | SE | *p* | Cohen's *d* | SE | *p* |
| Thalamus | -0.08 | 0.06 | 0.21 | -0.14 | 0.09 | 0.12 | - | - | - | -0.10 | 0.08 | 0.20 | -0.08 | 0.13 | 0.56 |
| Caudate | -0.04 | 0.06 | 0.59 | -0.02 | 0.09 | 0.83 | - | - | - | -0.02 | 0.08 | 0.81 | -0.16 | 0.13 | 0.23 |
| Putamen | 0.03 | 0.06 | 0.60 | 0.02 | 0.09 | 0.83 | - | - | - | 0.03 | 0.08 | 0.71 | -0.08 | 0.13 | 0.52 |
| Pallidum | 0.03 | 0.06 | 0.65 | -0.01 | 0.09 | 0.94 | - | - | - | 0.03 | 0.08 | 0.72 | -0.03 | 0.13 | 0.85 |
| Hippocampus | **-0.24** | **0.06** | **0.0003\*\*** | **-0.31** | **0.09** | **0.0005\*\*** | - | - | - | **-0.29** | **0.08** | **0.0004\*\*** | -0.24 | 0.13 | 0.063 |
| Amygdala | **-0.18** | **0.06** | **0.0058\*\*** | **-0.30** | **0.09** | **0.0008\*\*** | - | - | - | **-0.25** | **0.08** | **0.0024\*\*** | -0.13 | 0.13 | 0.31 |
| Accumbens | -0.05 | 0.06 | 0.42 | -0.11 | 0.09 | 0.23 | - | - | - | -0.07 | 0.08 | 0.40 | -0.06 | 0.13 | 0.62 |
| Lateral Ventricles | 0.08 | 0.06 | 0.20 | 0.12 | 0.09 | 0.19 | - | - | - | 0.12 | 0.08 | 0.13 | -0.01 | 0.13 | 0.96 |

\*\*Significant at Bonferroni corrected threshold for tests in 8 regions of interest $p \leq 0.0063$
\*Suggestive at $p \leq 0.05$

### Validation analyses

Validation analyses largely confirmed primary results. Please see **Supplementary Tables 6 and 7** for regional associations with dichotomized CD4+ counts (≤ 200 cells/mm^3^), and a harmonized dVL threshold (> 400 copies/mL), respectively. Results from the meta-analysis of CD4+ associations are reported in **Supplementary Table 8**, with corresponding Forest plots in **Supplementary Figures 4 and 5**

## Discussion

In one of the largest coordinated brain imaging studies of HIV+ individuals, we harmonized and pooled data from 1,044 individuals across six countries, and found associations between brain volumes and plasma markers used to monitor HIV infection. In the full group, brain volume associations specifically implicated the limbic system: lower CD4+ counts were associated with smaller hippocampal and thalamic volumes; a dVL was associated with smaller hippocampal and amygdala volumes. The limbic effects were largely driven by cART+ participants, the majority of whom were male. In the subset of cART-participants, however, smaller putamen volumes were associated with lower CD4+.

Plasma CD4+ T-cell count and VL are routinely assessed in HIV clinics around the world. These peripheral markers have been correlated with neuropsychological performance in HIV+ individuals,^1,13^ as well as *post mortem* brain tissue VL and pathology.^14,15^ Unfortunately, the relationship between plasma markers and MRI brain volume variation, assessed *in vivo*, has been inconsistent across studies. Differences between studies, including clinical and demographic differences, methodological variability, and/or insufficient power to estimate robust effect sizes, continue to complicate our understanding of the neuroanatomical and neurofunctional consequences of infection.

Brain signatures that generalize beyond individual studies are important to identify to establish neuroimaging biomarkers of HIV brain injury. Literature-based meta-analyses assess consistency of published findings, and offer initial insights into the generalizability of HIV-related findings. A recent such meta-analysis of 19 published studies, spanning almost three decades, found significantly lower total brain volumes and total gray matter volumes in HIV+ individuals compared to seronegative controls;^6^ however, no significant serostatus association with basal ganglia volume was found. In addition, effect sizes were found to be lower in more recent studies compared to older ones,^6^ which may suggest a diverging profile of HIV-affected neurocircuitry in more recent years.

The scope and extent of HIV-related neurocognitive impairment has evolved from the pre-cART era and some chronic HIV+ individuals in the cART era carry this legacy. For example, while the prevalence of neurocognitive deficits may be similar, reports have highlighted that learning deficits and poorer mental flexibility were more prominent features in cART-treated patients compared to pre-cART patients where deficits in motor and sustained attention and psychomotor slowing were more evident.^1,16^ Brain deficits may persist despite cART due to factors such as possible cART neurotoxicity, irreversible brain damage associated with advanced disease, reservoirs of ongoing low-grade viral replication and/or persistent immune activation in the CNS, vascular injury, and neurodegenerative processes that can occur with aging, consistent with the hippocampal findings presented here. ^3,4,6,17^

In the pre-cART era, HIV-associated dementia was characterized as a “subcortical dementia”. In line with reported *post mortem* neuropathological and viral protein distributions, neuroimaging studies of HIV+ individuals often showed smaller basal ganglia volumes.^18,19^ More recently, neuroimaging findings have extended beyond volume differences of the putamen, pallidum, nucleus accumbens and caudate. Serostatus related differences have been detected to varying degrees in the thalamus,^20^ amygdala,^21^ hippocampus,^22^ and even total gray matter,^6^ indicating that recent brain alterations may be more dispersed; this may also suggest greater heterogeneity in the causes of brain injury. Once a staple of HIV-related brain alterations, basal ganglia volume associations with CD4+ count or dVL were not detected in the full group of HIV+ participants assessed here, but only in the cART-subset. cART-participants were not necessarily cART naïve, but on average, their plasma markers may be more in line with those recorded before cART initiation. While current CD4+ counts were associated with altered limbic, and not basal ganglia, volumes in the full set of participants, it is possible that in these same individuals, nadir CD4+ counts —reflecting a history of severe immunosuppression prior to treatment— would be associated with basal ganglia volumes. The shift in HIV-infection from a fatal to chronic condition in the cART era appears to be accompanied by a shift in the profile of HIV-related brain abnormalities beyond the basal ganglia, frequently implicated in the pre-cART era, to limbic structures; this shift in subcortical signatures may be contributing to the growing range of neuropsychiatric and cognitive outcomes.

Males in our study may be more representative of the cART era: 90% of them were on cART compared to only 35% of females, most of whom were recruited from Thailand and South Africa –countries from which a substantial proportion of individuals eligible for treatment may not receive it. We found a similar pattern in the ranking of regional CD4+ count effect sizes between sexes, but regional effects were less consistent for dVL; females showed larger effect sizes in the caudate and putamen, while males showed larger effects in the amygdala and ventricles, further supporting a diverging cART era profile. As suggested in a US-based cohort of HIV+ women,^23^ demographic and social-cultural factors, which are themselves associated with many comorbid conditions, potentially mask the effect of HIV disease factors. Despite being the largest neuroimaging evaluation of HIV+ adult women, we did not detect any significant associations between plasma markers and brain volumes in females alone. This apparent difference in power may be due to any number of confounding factors. For example, comorbidities, including mental health conditions, may play a confounding role. Female-specific findings were noted in an international study of post-traumatic stress disorder (PTSD),^12^ where women with PTSD were found to be driving case-control differences in hippocampal volumes. Trauma, particularly related to intimate partner violence, is overwhelmingly common among HIV+ women,^24^ so it is possible that factors such as comorbid PTSD, may be confounding limbic associations in women. Women constitute 52% of all individuals aged 15 years and older living with HIV.^25^ Nevertheless, women are underrepresented in neuroHIV research, impeding the reliability and generalizability of findings. Despite evaluating over 300 HIV+ women, twice as many participants included in our study were male; only two of the 12 studies had recruited more than 40% women. A more extensive international effort assessing the neurological effects of HIV-infection in women is needed.

Effects in limbic structures, as seen here, have been detected in a wide-range of clinical conditions studied in similar large-scale international efforts from the ENIGMA consortium.^26^ Serious mental illnesses, such as depression or substance-use disorder, have a high prevalence in HIV+ individuals and may increase the risk of HIV transmission.^27^ As we assessed individuals with and without these comorbidities, limbic associations detected in this study are not likely simply a reflection of such comorbid neuropsychiatric conditions. However, viral-induced immunosuppression may also contribute to the risk of developing SMIs by targeting the same neurocircuitry implicated across these disorders.

The hippocampus, showed the largest effect size with both CD4+ and VL measures. In *post mortem* studies, hippocampal tissue shows some of the highest viral concentrations.^28^ Hippocampal neurons also show increased gliosis and HIV chemokine co-receptors and expression,^29^ and may be particularly susceptible to Tat-induced apoptosis.^30^ cART era pathological studies suggest a potential shift in HIV-related inflammation to the hippocampus and surrounding entorhinal cortex.^31^ Hippocampal atrophy is consistently reported across aging populations, and accelerated atrophy is a hallmark of neurodegenerative diseases such as Alzheimer’s disease (AD). Neuropathological hallmarks of healthy aging and AD, including elevated levels of phosphorylated tau and beta-amyloid deposits, have been detected in the hippocampus of cART treated HIV+ individuals.^32,33^ Common age and HIV-related pathological processes, such as inflammation and blood brain barrier impairment, may accelerate age-related neurodegenerative processes.^17^ Study participants ranged in age from 22 to 85 years, and while 1) the cART+ subgroup was older than the cART-group and 2) age was associated with reduced volumes in limbic structures, we did not detect any interaction between age and CD4+ count or dVL in the full group. A better understanding of chronic infection in the context of aging remains an important topic of research.

Literature based meta-analyses provide substantial insights into the reproducibility and consistency of published findings. However, they are inherently limited by the fact that effects are likely over-estimated due to publication biases substantially limiting the inclusion of studies with null findings. Effect sizes reported in this study are considered small to moderate, yet given the large, diverse sample, small effect sizes may not be clinically insignificant. Furthermore, methodological factors, including image analysis methods and statistical design cannot be harmonized in retrospective analyses. Here, we partially address these limitations by harmonizing image processing and statistical analyses. Nonetheless, several limitations and challenges remain. Our study focused only on associations in HIV+ individuals, without a direct comparison to seronegative controls. Serostatus comparisons may help elucidate the full extent of HIV-related brain deficits, as opposed to highlighting regions more selectively affected by the degree of immunosuppression and viral control. Most of the neuroHIV cohorts included in ENIGMA-HIV at the time of this study, however, recruited only HIV+ individuals. Plasma CD4+ count and VL were readily available across studies, but these plasma markers are not comprehensive assessments of the full systemic impact of HIV. Future efforts are needed to pool and harmonize additional immunological, cerebrovascular, metabolic, and inflammatory markers associated with infection. Identifying clinical factors that can be uniformly collected or interpreted across international studies remains challenging. For example, differences in treatment regimens with varying CNS penetration effectiveness (CPE) scores, duration of treatment, and standards of adherence that qualified a participant as cART+ at the time of scan, may vary from study to study. Despite such variations, our study identified consistent and robust brain volume associations with HIV VL and immunosuppression in a large and diverse study sample of HIV+ individuals from around the world.

This analysis demonstrates the feasibility and utility of a global collaborative initiative to understand the neurological signatures of HIV infection. We invite other neuroHIV researchers to join the ENIGMA-HIV consortium; with a greater collaborative effort, we will be able to assess factors that may modulate neurological outcomes –including cART treatment regimens, comorbidities, co-infections, substance use, socioeconomic, and demographic factors— as well as the functional implications of such structural brain differences, in well-powered analyses. Understanding the neurobiological changes that may contribute to neuropsychiatric and cognitive outcomes in HIV+ individuals is critical for identifying individuals at risk for neurological symptoms, driving novel treatments that may protect the CNS, and monitoring treatment response.

## Author contributions

TMN, JPF, PMT, RAC, RP, DJS, and NJ conceived and designed the analysis. TMN, JPF, JA, BMA, JB, BJB, LC, JRC, CRKC, LAC, TE, JF, VG, JH, JMHW, CHH, JH, JAJ, KJK, TK, HYL, ML, CLF, AJL, LM, BKN, BAN, XP, ECP, CMS, ADT, VV, MV, AJW, PMT, RAC, RP, DJS and NJ acquired, analyzed or interpreted the data. TMN and NJ wrote the first draft of the manuscript. LAC, RP, JPF, JA, BMA, JB, BJB, LC, JRC, CRKC, TE, JF, VG, JH, JMHW, CHH, JH, JAJ, KJK, TK, HYL, ML, CLF, AJL, LM, BKN, BAN, XP, ECP, CMS, ADT, VV, MV, AJW, PMT, RAC, and DJS critically revised the manuscript for important intellectual content. All authors approved the content of the final manuscript and agree to be accountable for the accuracy and integrity of all aspects of the work.

## Acknowledgements

Funding for the ENIGMA Center for Worldwide Medicine Imaging and Genomics was provided by the NIH Big Data to Knowledge Program (BD2K; U54 EB020403). Additional funding for this work was provided by NIH grants R01AG059874 (NJ), R01MH117601 (NJ), T32AG058507 (TMN and CRKC), 5T32MH073526 (CRKC), MH083553 (CHH), MH19535 (CHH), R01HL095135 (KJK), MH095661 (TK), UL1RR033176 (TK), UL1TR000124 (TK), MH19535 (TK), NS080655 (BAN), K01AA025306 (ECP), R01HL095135 (CMS), U54MD007584 (CMS), K23MH095661 (ADT), K23AG032872 (VV), R01NS061696 (VV), K24MH098759 (VV), K01AG050707 (AJW), P01AA019072 (RAC), R01MH074368 (RAC), P30 AI042853 (RAC), and by R01MH085604 (RP). This study was also supported by the SA Medical Research Council (DJS), NHMRC APP568746 and APP1045400 (LAC), and the European Research Council Advanced Grant MedYMA 2011-291080 (XP). We would like to thank all study participants and study groups.

