## Supplementary Appendix for "Smaller limbic structures are associated with greater immunosuppression in over 1000 HIV-infected adults across five continents: Findings from the ENIGMA-HIV Working Group"

### 1. Supplementary Data

#### 1.1 Inclusion and Exclusion Criteria, by Site

**Supplementary Table 1.** Inclusion and exclusion criteria for study participants.

| Site name | Inclusion criteria | Exclusion criteria | Reference |
|---|---|---|---|
| **HIVNC Consortium (7 Sites), United States** | HIV positive; Age > 18 years; On stable cART ≥ 12 weeks; Nadir CD4 < 200 cells/mm$^3$ | Major psychiatric illness; Confounding neurological disorders; Brain infection other than HIV; Hepatic dysfunction; Diabetes mellitus; Active substance abuse or related medical complications within 6 months of study | Nir et al. 2019[1] |
| **University of Hawaii, United States** | HIV positive; Age > 40 years; On stable cART ≥ 3 months | Uncontrolled major affective disorder; Active psychosis; Loss of consciousness > 5 min; Pregnant or breastfeeding; Any past or present confounding condition (e.g. CNS infection, TBI, stroke, substance abuse) | Kallianpur et al. 2016[2] |
| **University of California, San Francisco, United States** | Age > 60 years | Learning disabilities; Major psychiatric or neurological illness; Current or past brain infection; Major systemic illness or head injury | Clifford et al. 2017[3] |
| **Brown University, United States** | Age > 18 years<br><br>Note: Some HIV+ participants presented with Hepatitis C | History of head injury with loss of consciousness > 10 min; Neurological conditions including dementia unrelated to HIV, seizure disorder, stroke, and opportunistic infection of the brain; Severe psychiatric illness that may impact brain function (e.g., schizophrenia); Diagnosis of alcohol or substance abuse or dependence based within 6 months prior to neuroimaging | Gongvatana et al. 2014[4] |
| **University of California, Los Angeles, United States (Hinkin)** | Age > 18 years | Neurological, psychiatric, and medical confounds (e.g., history of seizure disorder or other neurologic disorder; history of concussion or TBI; history of Axis I psychiatric disorder or current substance use disorder; current prescription for psychotropic medication, except for anxiolytics and antidepressants; current substance dependence or methamphetamine use); Comorbid CNS infection (e.g. Hepatitis C); HIV-associated CNS opportunistic infection (e.g. toxoplasmosis) or neoplasm; MRI contraindications | Kuhn et al. 2017[5] |
| **University of California, Los Angeles, United States (Thames)** | Age > 18 years; Chronic HIV infection (mean duration of 12 years); On stable cART ≥ 3 months; Clinically stable (as indicated by CD4+ cells/mm$^3$); Score ≥ 26 on Mini-Mental Status Exam | Current abuse of cocaine/amphetamines; Past stimulant abuse/dependence; Current/past diagnosis of psychotic spectrum disorder; CNS confounds (e.g., HIV-associated opportunistic infections or neuro-syphilis); Hepatitis C coinfection; Major head injury; MRI contraindication | Thames et al. 2018[6] |
| **University of New South Wales, Australia (Brew)** | Age > 18 years; On stable cART ≥ 12 months | Contraindication to MRI; Comorbid CNS infection, tumour or stroke | Changanti et al 2017[7] |
| **University of New South Wales, Australia (Cysique)** | Age > 45 years; On stable cART ≥ 6 months; nadir CD4 ≤ 350 cells/mm$^3$; Known HIV duration ≥ 5 years; No active opportunistic disease<br><br>Note: HIV+ participants all male | MRI contraindication; History of neurological disorders predating HIV diagnosis (e.g., epilepsy, traumatic brain injury, Parkinson's Disease, Multiple Sclerosis, Alzheimer's disease, or Vascular Dementia); Psychiatric disorders on the psychotic axis (e.g., schizophrenia); Current substance use disorders (within 12 months of study enrolment; recreational use of marijuana was not set as a criterion for exclusion because it would exclude a large number of HIV+ individuals); Participants not excluded on the basis of current depressive symptoms; Hepatitis C status was recorded from the participants' medical records and participants were included only if successfully treated and/or inactive (N=2) | Nichols et al. 2019[8] |
| **SEARCH 011 Consortium, Thailand** | Age > 18 years; cART-naïve; CD4 < 350 cells/mm$^3$ or symptomatic HIV | Head injury; Current substance abuse; Acute concurrent illness; Neurologic or psychiatric conditions; Learning disabilities; Positive hepatitis C screen; Contraindications for MRI; Significant laboratory abnormalities (e.g. creatinine, ALT, hemoglobin) | Heaps et al. 2015[9] |

| | | | |
|---|---|---|---|
| **University of Cape Town, South Africa** | Age 18 to 50 years to avoid CNS complications associated with neurodevelopment and advanced age; at least 5 years of formal education; cART initiation within 3 months of enrollment; Xhosa as the primary language<br><br>Note: Predominantly female; 100% detectable viral load | Major psychiatric conditions (schizophrenia, bipolar disorder, post-traumatic stress disorder, etc.); Neurological disease that could affect brain integrity (e.g., multiple sclerosis); CDC stage A; Opportunistic infections of the CNS (e.g., cytomegalovirus encephalitis, cryptococcal meningitis, toxo- plasma encephalitis); Lifetime history of head injury resulting in loss of consciousness >30 min; Current substance use disorder as determined by the Mini- International Neuropsychiatric Interview Plus | Paul et al. 2017[10] |
| **Nice University, France** | HIV positive; age > 18 years | CNS opportunistic infections; Change in psychotropic therapy < 3 weeks; Any neurological history | Vassallo et al. 2015[11] |
| **University of Novi Sad, Serbia** | Age > 18 yrs; On stable cART ≥ 12 months; Nadir CD4+ ≤ 250 cells/mm$^3$ | A history of neurological disorders predating HIV (e.g., epilepsy, brain tumor, TBI, Parkinson's Disease, Multiple Sclerosis, Alzheimer's disease, or Vascular Dementia); Psychiatric disorders on the psychotic axis (e.g., schizophrenia); Cardiovascular diseases (hypertension, chronic occlusive carotid disease, ischemic vascular dementia); History of or current substance use disorders; Hepatitis C (HCV) or Hepatitis B positive | Boban et al. 2017[12] |

#### 1.2 MRI Parameters, by Site

**Supplementary Table 2.** T1-weighted MRI acquisition parameters

| Site | Scanner | Acquisition parameters |
|---|---|---|
| **HIVNC Consortium, United States** | 1.5T GE Signa, Siemens Symphony/Sonata | GE SPGR; TR=2-23 ms; TE=3-9 ms; flip angle=30°; matrix size = 256 × 128; voxel size=1x1x1.2-1.5 mm<br>Siemens: TR=20-24 ms; TE=10.1 ms; flip angle = 30°; matrix size = 256 × 192; voxel size=1x1x1.2-1.3 mm; |
| **University of Hawaii, United States** | 3.T Phillips Achieva | 3D TFE; TR=6.9 ms; TE = 3.2 ms; flip angle = 8°; FOV = 256mm; voxel size=1x1x1.2 mm |
| **University of California, San Francisco, United States** | 3T Siemens TrioTim | MP-RAGE; TR=2300 ms; TE=2.98 ms; flip angle=9°; FOV = 256mm; 160 slices; 240×256 matrix; voxel size=1x1x1 mm |
| **Brown University, United States** | 3T Siemens TrioTim | MP-RAGE; TR=2250 ms; TE=3.06 ms; flip angle=9°; FOV = 220 mm; matrix = 256×256, slice thickness = 0.86 mm |
| **University of California, Los Angeles, United States (Hinken)** | 3T Siemens TrioTim | MP-RAGE; TR=2200 ms; TE=2.2 ms; matrix size = 256 × 256; FOV 240 mm; 176 slices; slice thickness = 1 mm |
| **University of California, Los Angeles, United States (Thames)** | 3T Siemens TrioTim | MP-RAGE; TR = 450.0 ms; TE = 10.0 ms; flip angle: 8°; FOV 256 mm; matrix =256x219; voxel size =1.0 × 0.94 × 0.94 mm |
| **University of New South Wales, Australia (Brew)** | 3T Philips Achieva | 3D TFE; TR=5.3 ms; TE=2.4ms; flip angle = 18°, 256x256 matrix; 180 slices; voxel size=1x1x1 mm |
| **University of New South Wales, Australia (Cysique)** | 3T Philips Achieva | 3D TFE; TR= 6.39ms; TE=2.9 ms; flip angle: 8°; FOV 256 mm; 190 slices, voxel size=1x1x1 mm |
| **SEARCH 011 Consortium, Thailand** | 3T Siemens Allegra | MP-RAGE; TR=2400 ms; TE=2.38 ms; flip angle=8°; 162 slices; voxel size=1x1x1 mm |
| **University of Cape Town, South Africa** | 3T Siemens Allegra | MP-RAGE; TR=2400 ms; TE=2.38 ms; TI=1000 ms; flip angle=8°; 162 slices; voxel size=1x1x1 mm |
| **Nice University, France** | 1.5T GE Signa | SPGR; TR =12.4 ms, TE = 5.2 ms, flip angle = 18°; FOV = 240 mm; 256x256 matrix; voxel size=0.6 x 0.6 x 0.6 mm |
| **University of Novi Sad, Serbia** | 3T Siemens TrioTim | MP-RAGE; TR = 2300 ms; TE=2.97 ms; FOV 256 mm; voxel size=1 x 1 x 1 mm |

##### 1.3 Viral Load Thresholds, by Site

**Supplementary Table 3.** Approximate detectable plasma RNA viral load threshold by site.

| Site | Detectable Viral Load Threshold (copies/mL) |
|---|---|
| **HIVNC Consortium (7 Sites), United States** | 50-400 |
| **Site 1: University of California, San Diego** | 50 |
| **Site 2: Harbor UCLA Medical Center** | 75 |
| **Site 3: Stanford University** | 50 |
| **Site 4: Colorado** | 50 |
| **Site 5: Pittsburgh** | 50 |
| **Site 6: Rochester University** | 50 |
| **Site 7: University of California, Los Angeles** | 400 |
| **University of Hawaii, United States** | 50 |
| **University of California, San Francisco, United States** | 50 |
| **Brown University, United States** | 75 |
| **University of California, Los Angeles, United States (Hinken)** | 48 |
| **University of California, Los Angeles, United States (Thames)** | 19 |
| **University of New South Wales, Australia (Brew)** | 20 |
| **University of New South Wales, Australia (Cysique)** | 50 |
| **SEARCH 011 Study, Thailand** | 40 |
| **University of Cape Town, South Africa** | 50 |
| **Nice University, France** | 40 |
| **University of Novi Sad, Serbia** | 50 |

#### 2. Supplementary Methods and Results

##### 2.1 Age Mega-Analysis

To ensure sufficient power to detect associations between brain measures and variables of interest from data pooled across studies, a preliminary analysis tested for associations between age and eight brain volumes. Random effects multiple linear regressions were performed. Two fixed-effects covariates were included in the model: sex and total intracranial volume (ICV) to adjust for variability in head size. To account for scanner effects, the data collection site was used as the random-effects grouping variable; there were 18 sites in all. Statistical analyses were conducted with the 'nlme' package in R (version 3.2.3). Effect sizes were estimated using the *r*-value (partial correlation coefficients) after accounting for all covariates. Significance was determined using the Bonferroni correction threshold ($p \leq 0.0063$).

Across the entire sample (N=1,044), older age was associated with smaller volumes of all subcortical structures and larger ventricular volumes (*r*-value range: 0.13-0.29), consistent with studies in seronegative adults.[13] Sex was a significant predictor in models for associations between age and the thalamus, putamen, pallidum, hippocampus, and amygdala ($p \leq 0.0063$). Age effects in the subset of cART+ participants and males mirrored that of the full group. cART- participants showed significant negative associations between age and caudate, putamen, pallidum and nucleus accumbens volumes. Females showed significant negative associations between age and thalamus, putamen, pallidum, nucleus accumbens, and lateral ventricle volumes. No significant age by sex interaction was detected.

**Supplementary Table 4.** *R*-values (partial correlation coefficients), *b*-values (unstandardized regression slopes reflecting change in volume ($mm^3$) for every year of age), standard errors (SE), and *p*-values from associations between age and regional brain volumes across all 1,044 HIV+ participants, and separately in the subset of 756 cART+ participants, 288 cART- participants, 734 males, and 310 females.

| ROI | Total (n=1,044) | | | | cART+ (n=756) | | | | cART- (n=288) | | | | Male (n=734) | | | | Female (n=310) | | | |
|---|---|---|---|---|---|---|---|---|---|---|---|---|---|---|---|---|---|---|---|---|
| | *r* | *b* | SE | *p* | *r* | *b* | SE | *p* | *r* | *b* | SE | *p* | *r* | *b* | SE | *p* | *r* | *b* | SE | *p* |
| Thalamus | -0.24 | -25.26 | 3.16 | 3.78E-15** | -0.29 | -31.37 | 3.82 | 9.90E-16** | -0.13 | -14.33 | 6.77 | 0.035* | -0.34 | -28.62 | 2.96 | 7.08E-21** | -0.21 | -20.22 | 5.61 | 0.0004** |
| Caudate | -0.13 | -7.31 | 1.75 | 3.12E-5** | -0.085 | -5.19 | 2.24 | 2.07E-2** | -0.19 | -11.58 | 3.60 | 0.0014** | -0.15 | -6.96 | 1.68 | 3.68E-5** | -0.16 | -8.55 | 3.12 | 0.0065* |
| Putamen | -0.29 | -24.29 | 2.51 | 2.73E-21** | -0.27 | -25.07 | 3.31 | 1.15E-13** | -0.26 | -20.98 | 4.69 | 1.11E-5** | -0.33 | -24.01 | 2.54 | 4.35E-20** | -0.36 | -27.17 | 4.06 | 1.15E-10** |
| Pallidum | -0.21 | -5.99 | 0.88 | 1.76E-11** | -0.17 | -5.64 | 1.17 | 1.80E-6** | -0.23 | -6.60 | 1.68 | 1.06E-4** | -0.22 | -5.20 | 0.87 | 3.29E-9** | -0.29 | -7.75 | 1.51 | 5.51E-7** |
| Hippocampus | -0.16 | -9.01 | 1.70 | 1.34E-7** | -0.14 | -8.42 | 2.17 | 1.17E-4** | -0.13 | -7.61 | 3.53 | 0.032* | -0.24 | -10.84 | 1.66 | 1.20E-10** | -0.13 | -6.30 | 2.93 | 0.032* |
| Amygdala | -0.15 | -3.77 | 0.80 | 3.15E-6** | -0.12 | -3.27 | 1.04 | 1.72E-3** | -0.11 | -2.79 | 1.54 | 0.071 | -0.20 | -4.33 | 0.80 | 7.12E-8** | -0.14 | -3.38 | 1.39 | 0.016* |
| Accumbens | -0.25 | -3.19 | 0.39 | 1.15E-15** | -0.21 | -3.02 | 0.51 | 5.49E-9** | -0.21 | -2.77 | 0.79 | 0.0005** | -0.26 | -2.84 | 0.40 | 3.54E-12** | -0.31 | -3.75 | 0.67 | 4.35E-8** |
| Lateral Ventricles | 0.20 | 326.47 | 49.65 | 7.74E-11** | 0.19 | 352.09 | 68.87 | 4.07E-7** | 0.15 | 199.73 | 80.02 | 0.013* | 0.27 | 383.52 | 51.63 | 3.16E-13** | 0.17 | 193.75 | 64.59 | 0.0029** |

**Significant at Bonferroni corrected threshold for tests in 8 regions of interest, $p \leq 0.0063$

*Suggestive at $p \leq 0.05$

#### 2.2 Age Meta-Analysis

To confirm findings from analyses where data were pooled (mega-analyses), we performed a meta-analyses, where effects were found separately for each participating study and effect sizes were then aggregated using an inverse-variance weighted meta-analysis. Multiple linear regressions were performed covarying for sex and ICV. Resulting effect sizes (*b*-values) across each of the 18 scanning sites were then aggregated using an inverse-variance weighted fixed-effects model from the 'metafor' package in R. Forest plots were created to compare individual site effects and the aggregated effects. Associations between regional brain volumes and age *meta-analyzed* across 18 sites are consistent with pooled findings.

**Supplementary Table 5.** We report *r*-values (partial correlation coefficients), *b*-values (reflecting change in $mm^3$ volume for every year of age), standard errors (SE), heterogeneity scores ($I^2$) indicating the percentage of the total variance in effect size explained by heterogeneity across sites, and *p*-values from associations between regional brain volumes and age *meta-analyzed* across 18 sites. Forest plots (**Supplementary Figures 1 and 2**) show site-specific effects, compared with pooled mega, and meta-analytical effect sizes.

| ROI | Meta-Analysis | | | | |
|---|---|---|---|---|---|
| | *r* | *b* | SE | $I^2$ | *p* |
| **Thalamus** | **-0.27** | **-27.45** | **3.04** | **20.16** | **1.79E-19**** |
| **Caudate** | **-0.097** | **-5.93** | **1.88** | **35.76** | **0.0016**** |
| **Putamen** | **-0.25** | **-22.87** | **2.72** | **0** | **4.72E-17**** |
| **Pallidum** | **-0.16** | **-4.77** | **0.92** | **59.19** | **2.22E-7**** |
| **Hippocampus** | **-0.17** | **-9.81** | **1.81** | **49.53** | **6.01E-8**** |
| **Amygdala** | **-0.11** | **-3.01** | **0.82** | **24.34** | **0.0002**** |
| **Accumbens** | **-0.18** | **-2.32** | **0.40** | **0** | **4.77E-9**** |
| **Lateral Ventricles** | **0.21** | **311.88** | **44.60** | **21.99** | **2.69E-12**** |

**Significant at Bonferroni corrected threshold for tests in 8 regions of interest, $p \leq 0.0063$

#### 2.3 Forest Plots for Effects of Age

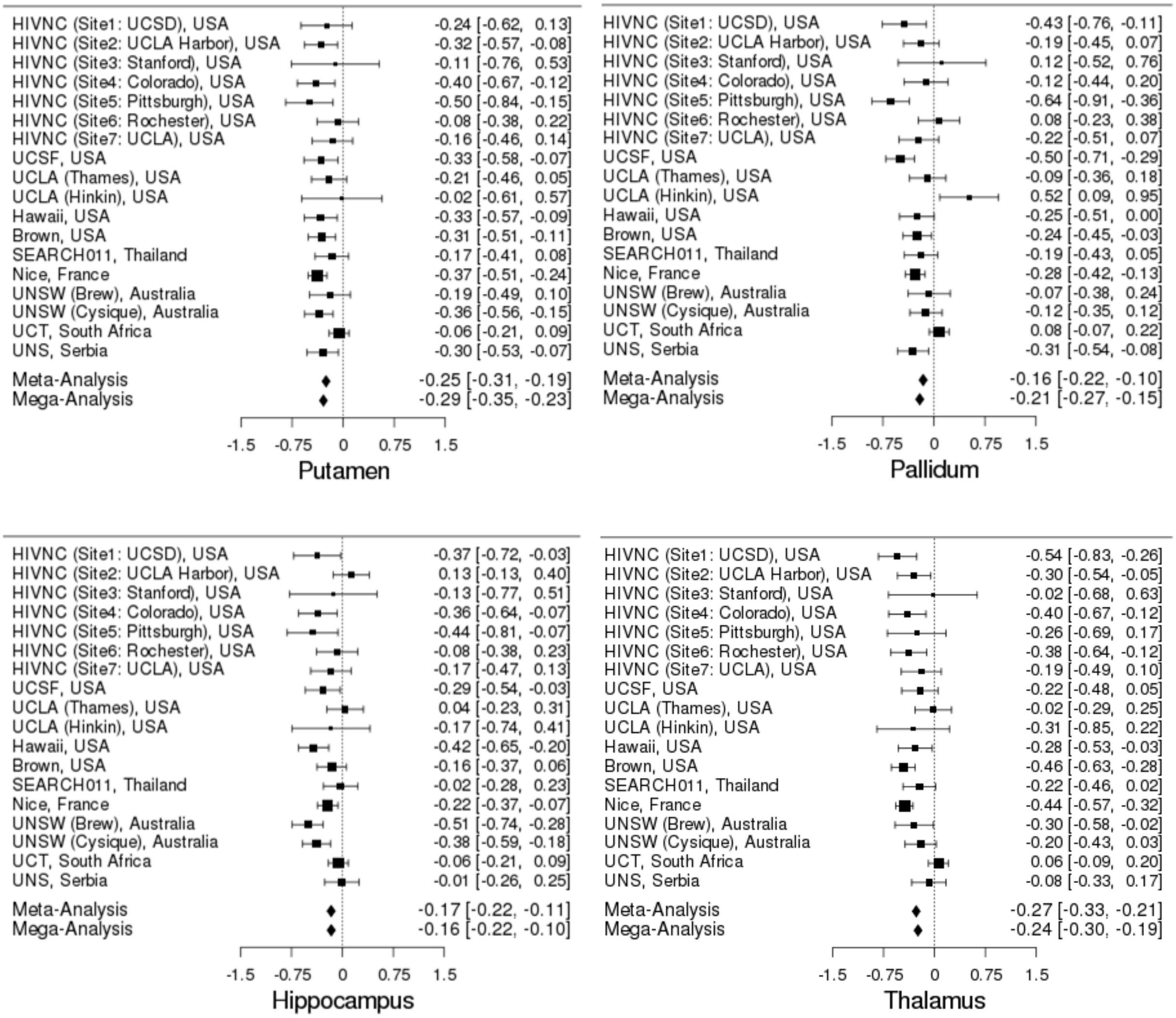

**Supplementary Figure 1.** Forest plot of effect sizes (*r*-values and 95% confidence intervals) for associations between age and putamen, pallidum, hippocampal or thalamic volumes across all 18 sites.

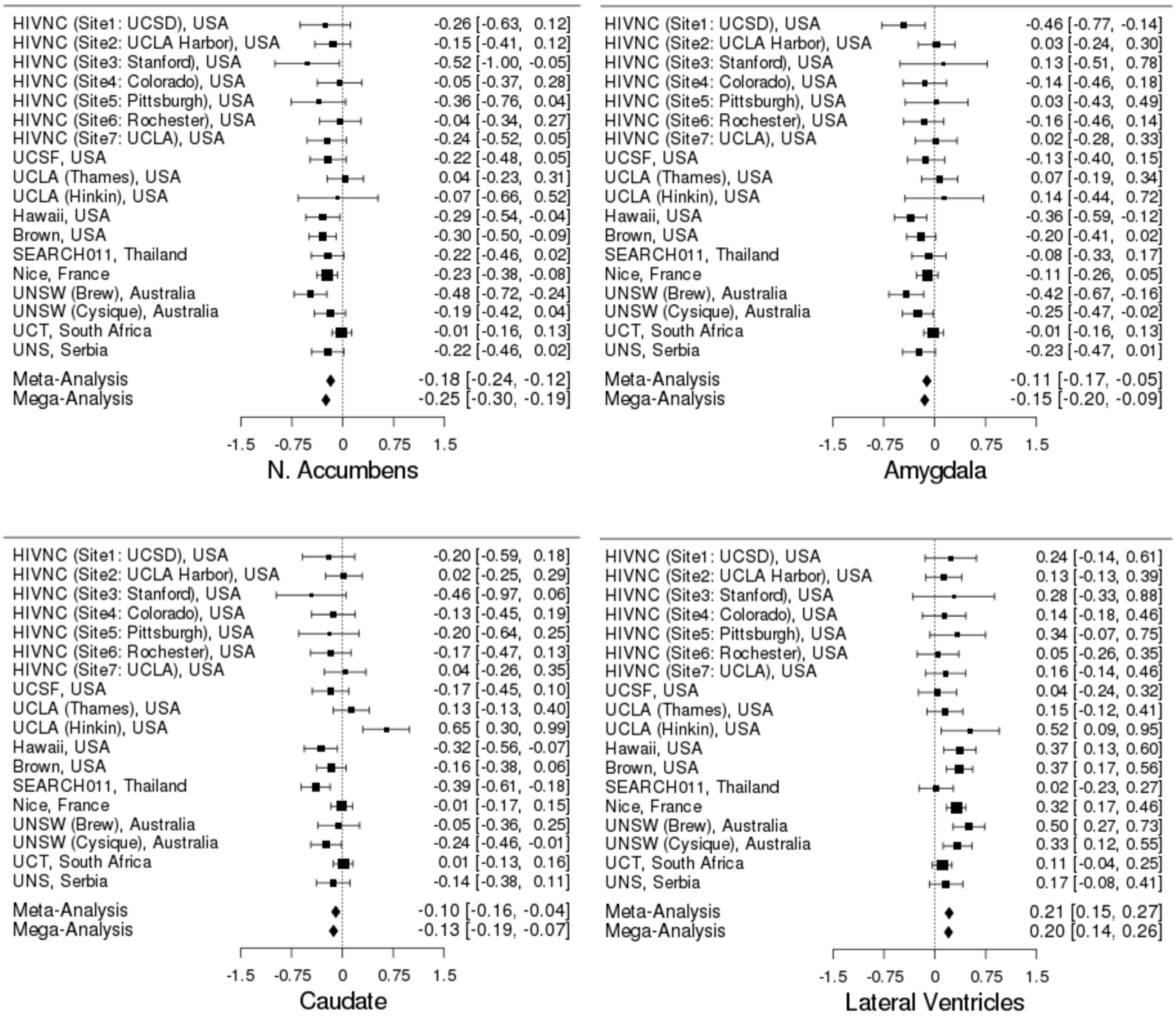

**Supplementary Figure 2.** Forest plot of effect sizes (*r*-values and 95% confidence intervals) for associations between age and accumbens, amygdalar, caudate or ventricular volumes across 18 sites. As expected, older age was associated with smaller volumes of all subcortical structures and larger ventricular volumes.

##### 2.4 Effect Sizes in Males Compared to Females

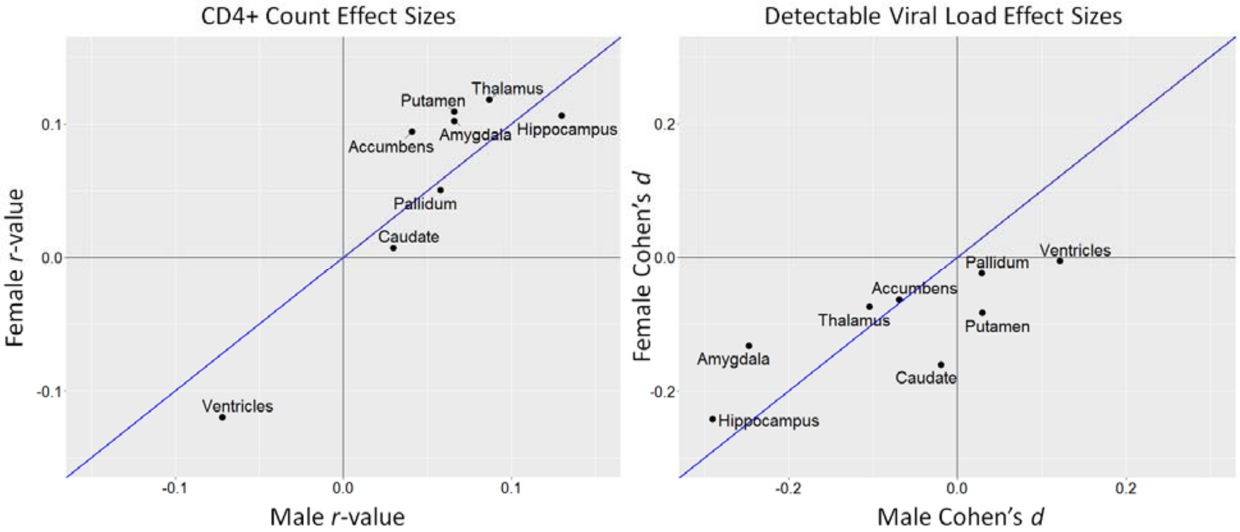

**Supplementary Figure 3.** Effect sizes in males compared to females for each ROI were highly correlated (CD4+: Pearson's correlation $r = 0.91$, $p = 0.002$; dVL: $r = 0.79$, $p = 0.02$).

##### 2.5 Validation Analyses

Three sets of validation analyses were performed: 1) dichotomizing CD4+ count based on the AIDS-defining threshold of 200 cells/mm$^3$; 2) defining a common dVL threshold across sites (400 copies/mL, the highest detection limit for any site); 3) comparing primary pooled statistical analyses to the meta-analytic framework used in other ENIGMA studies.[14] To confirm findings from HIV plasma marker analyses where data were pooled (mega-analyses), we performed a meta-analyses, where effects were found separately for each participating study and effect sizes were then aggregated using an inverse-variance weighted meta-analysis. Multiple linear regressions were performed covarying for age, sex, age-by-sex interaction, and ICV. Resulting effect sizes (*b*-values) across each of the 18 scanning sites were then aggregated using an inverse-variance weighted fixed-effects model from the 'metafor' package in R. Forest plots were created to compare individual site effects and the aggregated effects.

Suggestive associations were detected between brain volumes and an AIDS-defining immunosuppression status (CD4+ ≤ 200 cells/mm$^3$; **Supplementary Table 6**). dVL > 400 copies/mL was significantly associated with smaller hippocampal volumes ($d = -0.23$; $p = 0.0007$; **Supplementary Table 7**). When CD4+ count associations were meta-analyzed, findings were consistent regardless of study design, see **Supplementary Table 8** and **Supplementary Figures 4 and 5** for Forest plots. The distribution of the number of individuals with dVL was uneven across sites (see **Table 1**), preventing a comparable meta-analysis for dVL. One finding from the validation analyses was not detected in the main analysis: females with CD4+ ≤ 200 cells/mm$^3$ had significantly smaller nucleus accumbens volumes ($d = -0.37$; $p = 0.0035$).

##### 2.5.1 Dichotomized CD4+ Count Threshold Analysis

**Supplementary Table 6.** Cohen's *d* effect sizes for associations between regional brain volumes and an AIDS-defining immunosuppression status (CD4+ count ≤ 200 cells/mm$^3$) across 1,044 HIV+ individuals, and separately in the subset of 756 cART+ participants, 288 cART- participants, 734 males, and 310 females.

| ROI | Total (n=1044) | | | cART+ (n=756) | | | cART- (n=288) | | | Male (n=734) | | | Female (n=310) | | |
|---|---|---|---|---|---|---|---|---|---|---|---|---|---|---|---|
| | Cohen's *d* | SE | *p* | Cohen's *d* | SE | *p* | Cohen's *d* | SE | *p* | Cohen's *d* | SE | *p* | Cohen's *d* | SE | *p* |
| **Thalamus** | -0.092 | 0.078 | 0.24 | -0.11 | 0.12 | 0.34 | -0.18 | 0.12 | 0.14 | 0.020 | 0.11 | 0.85 | -0.25 | 0.12 | 0.050* |
| **Caudate** | -0.099 | 0.078 | 0.29 | -0.17 | 0.12 | 0.15 | -0.060 | 0.12 | 0.62 | -0.12 | 0.11 | 0.26 | -0.072 | 0.12 | 0.57 |
| **Putamen** | -0.20 | 0.078 | 0.011* | -0.16 | 0.12 | 0.18 | **-0.39** | **0.12** | **0.0014**** | -0.22 | 0.11 | 0.040* | -0.22 | 0.12 | 0.074 |
| **Pallidum** | -0.17 | 0.078 | 0.036* | -0.16 | 0.12 | 0.19 | -0.25 | 0.12 | 0.040* | -0.15 | 0.11 | 0.16 | -0.20 | 0.12 | 0.11 |
| **Hippocampus** | -0.18 | 0.078 | 0.024* | -0.062 | 0.12 | 0.60 | -0.24 | 0.12 | 0.049* | -0.21 | 0.11 | 0.050* | -0.17 | 0.12 | 0.19 |
| **Amygdala** | -0.15 | 0.078 | 0.062 | -0.11 | 0.12 | 0.38 | -0.19 | 0.12 | 0.12 | -0.098 | 0.11 | 0.37 | -0.23 | 0.12 | 0.073 |
| **Accumbens** | -0.14 | 0.078 | 0.069 | -0.11 | 0.12 | 0.34 | -0.27 | 0.12 | 0.028* | -0.030 | 0.11 | 0.78 | **-0.37** | **0.12** | **0.0035**** |
| **Lateral Ventricles** | -0.002 | 0.078 | 0.98 | -0.17 | 0.12 | 0.15 | 0.13 | 0.12 | 0.29 | -0.11 | 0.11 | 0.30 | 0.28 | 0.12 | 0.028* |
| **% with CD4+ Count < 200 cells/mm$^3$** | 19.35% | | | 12.33% | | | 42.01% | | | 13.76 % | | | 32.58% | | |

**Significant at Bonferroni corrected threshold for tests in 8 regions of interest, $p \leq 0.0063$

*Suggestive at $p \leq 0.05$

##### 2.5.2 Harmonized Viral Load Threshold Analysis

**Supplementary Table 7.** Cohen's *d* effect sizes for associations between regional brain volumes and dVL > 400 copies/mL across 1,006 HIV+ individuals, and separately in the subset of 752 cART+ participants, 724 males, and 282 females. We did not assess dVL in those off treatment due to the limited number of individuals in this subgroup with undetectable VL (n=20).

| ROI | Total (n=1,006) | | | cART+ (n=752) | | | cART- (n=254) | | | Male (n=724) | | | Female (n=282) | | |
|---|---|---|---|---|---|---|---|---|---|---|---|---|---|---|---|
| | Cohen's *d* | SE | *p* | Cohen's *d* | SE | *p* | Cohen's *d* | SE | *p* | Cohen's *d* | SE | *p* | Cohen's *d* | SE | *p* |
| **Thalamus** | -0.12 | 0.067 | 0.078 | -0.21 | 0.11 | 0.065 | -- | -- | -- | -0.16 | 0.090 | 0.084 | -0.037 | 0.12 | 0.77 |
| **Caudate** | -0.011 | 0.067 | 0.88 | 0.10 | 0.11 | 0.39 | -- | -- | -- | 0.056 | 0.090 | 0.54 | -0.14 | 0.12 | 0.26 |
| **Putamen** | 0.037 | 0.067 | 0.58 | 0.054 | 0.11 | 0.63 | -- | -- | -- | 0.035 | 0.090 | 0.70 | 0.022 | 0.12 | 0.86 |
| **Pallidum** | 0.031 | 0.067 | 0.65 | 0.069 | 0.11 | 0.54 | -- | -- | -- | 0.094 | 0.090 | 0.30 | -0.11 | 0.12 | 0.39 |
| **Hippocampus** | **-0.23** | **0.067** | **0.0007**** | **-0.39** | **0.11** | **0.0005**** | -- | -- | -- | **-0.34** | **0.091** | **0.0003**** | -0.075 | 0.12 | 0.55 |
| **Amygdala** | -0.083 | 0.067 | 0.22 | -0.19 | 0.11 | 0.090 | -- | -- | -- | -0.11 | 0.090 | 0.22 | -0.056 | 0.12 | 0.66 |
| **Accumbens** | -0.054 | 0.067 | 0.43 | -0.16 | 0.11 | 0.16 | -- | -- | -- | -0.11 | 0.090 | 0.23 | 0.074 | 0.12 | 0.56 |
| **Lateral Ventricles** | 0.11 | 0.067 | 0.11 | 0.23 | 0.11 | 0.037* | -- | -- | -- | 0.21 | 0.091 | 0.022* | -0.030 | 0.12 | 0.81 |
| **% with Viral Load > 400 copies/mL** | 32.70% | | | 12.63% | | | 92.13% | | | 21.55% | | | 61.35% | | |

**Significant at Bonferroni corrected threshold for tests in 8 regions of interest, $p \leq 0.0063$

*Suggestive at $p \leq 0.05$

##### 2.5.3 CD4+ Count Meta-Analysis

**Supplementary Table 8.** Associations between regional brain volumes and current CD4+ count *meta-analyzed* across 18 sites reflect similar findings to pooled findings. We report *r*-values (partial correlation coefficients), *b*-values (unstandardized regression slopes reflecting change in volume ($mm^3$) for every 100 cells/$mm^3$ change in CD4+ count), standard errors (SE), heterogeneity scores ($I^2$) indicating the percentage of the total variance in effect size explained by heterogeneity between sites, and *p*-values.

| ROI | Meta-Analysis | | | | |
|---|---|---|---|---|---|
| | *r* | *b* | SE | $I^2$ | *p* |
| **Thalamus** | **0.091** | **25.24** | **8.56** | **0.00** | **0.0031**** |
| **Caudate** | 0.010 | 1.73 | 5.43 | 2.99 | 0.75 |
| **Putamen** | 0.076 | 19.07 | 7.71 | 12.26 | 0.013* |
| **Pallidum** | 0.079 | 6.76 | 2.63 | 5.54 | 0.010* |
| **Hippocampus** | **0.11** | **17.21** | **5.02** | **32.24** | **0.0006**** |
| **Amygdala** | 0.071 | 5.25 | 2.27 | 0.000 | 0.021* |
| **Accumbens** | 0.057 | 2.19 | 1.19 | 32.24 | 0.065 |
| **Lateral Ventricles** | -0.082 | -341.09 | 128.34 | 0.00 | 0.0079* |

**Significant at Bonferroni corrected threshold for tests in 8 regions of interest, $p \leq 0.0063$
*Suggestive at $p \leq 0.05$

##### 2.5.4 CD4+ Count Forest Plots

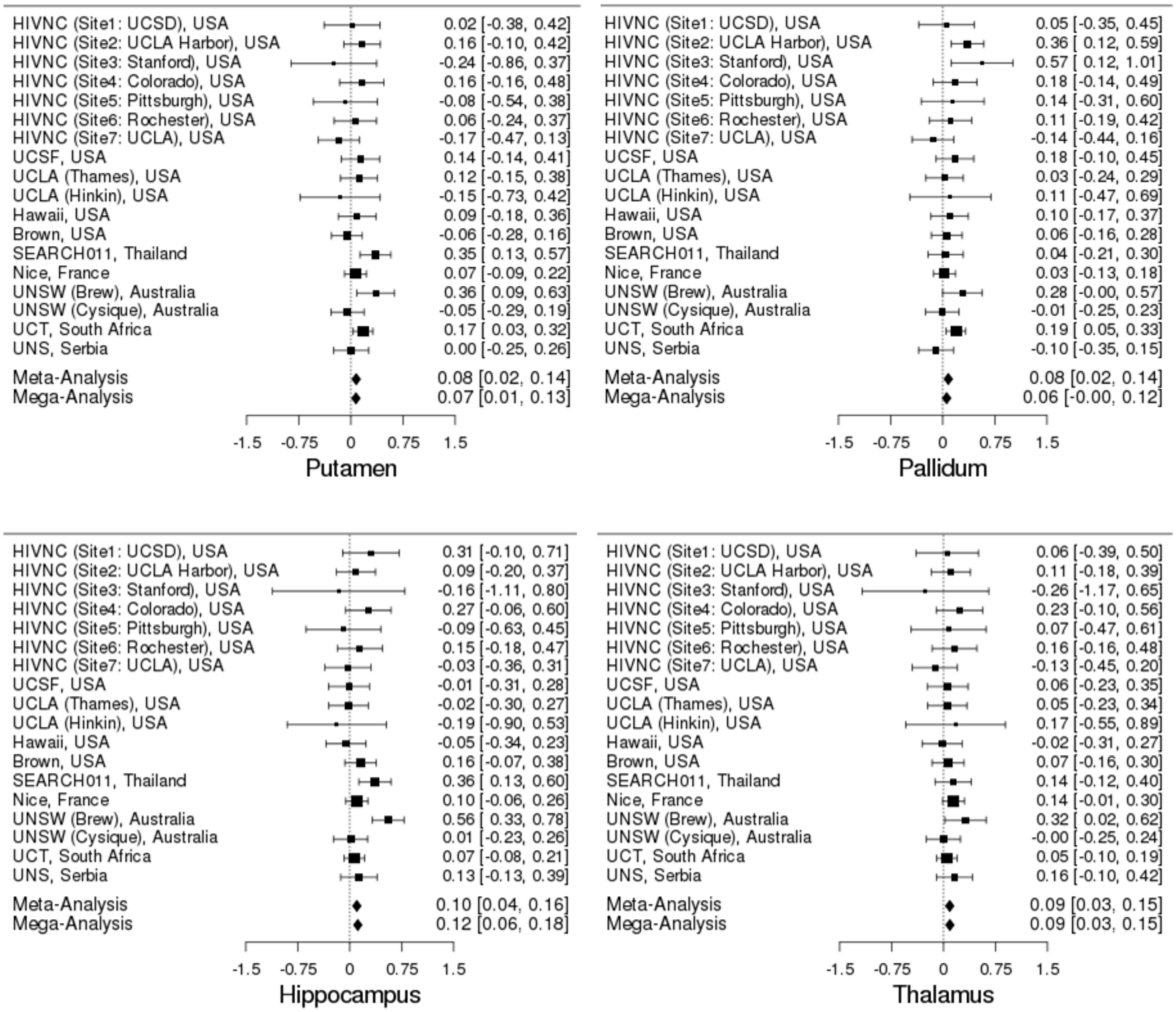

**Supplementary Figure 4.** Forest plot of effect sizes (*r*-values and 95% confidence intervals) for associations between CD4+ count and putamen, pallidum, hippocampal or thalamic volumes across 18 sites.

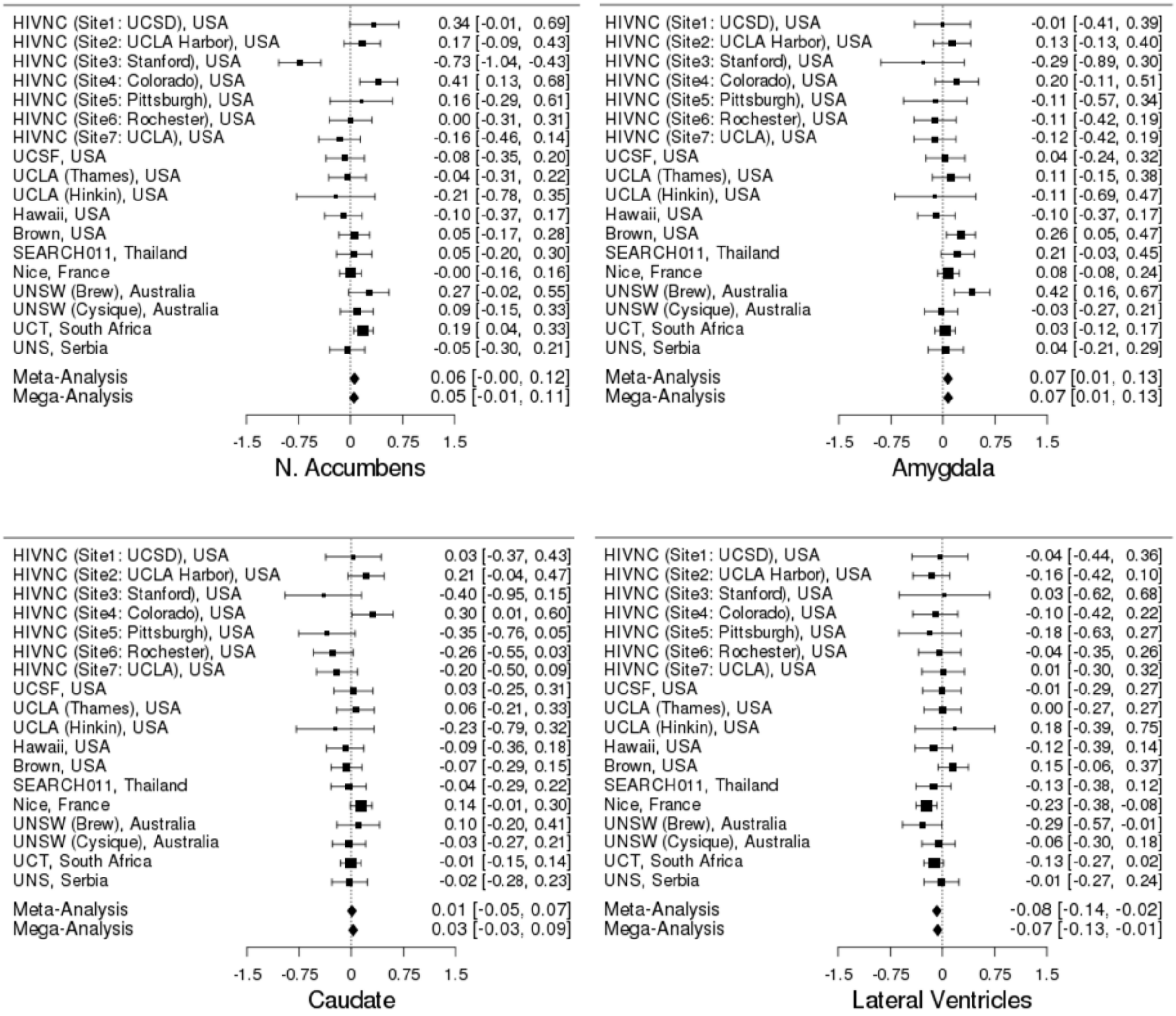

**Supplementary Figure 5.** Forest plot of effect sizes (*r*-values and 95% confidence intervals) for associations between CD4+ count and accumbens, amygdalar, caudate or ventricular volumes across 18 sites.
